# Epigenetic study of the long-term effects of Gulf War illness

**DOI:** 10.1101/2024.11.29.626040

**Authors:** B. C. Jones, J. P. O’Callaghan, D. G. Ashbrook, L. Lu, P. Prins, W. Zhao, K Mozhui

## Abstract

Gulf war illness is a chronic multisymptom disorder that affects as many as many as 25-35% of the military personnel who were sent to the Persian Gulf war in 1991. The illness has many debilitating symptoms, including cognitive problems, gastrointestinal symptoms, and musculoskeletal pain. Those so afflicted have been sick for more than 30 years and, therefore, it has become imperative to understand the etiology and then produce treatments to ease the symptoms. We hypothesized that the length of the disease was reflected in epigenetic modification of possibly several genes related to the symptoms. We subjected male and female mice from 11 BXD strains to combined corticosterone and the sarin surrogate, diisopropylfluorophosphate, to emulate the physiological stress of war and the potential exposures to organophosphate pesticides and nerve agent in theater. Three hundred days after treatment, we analyzed the animals’ DNA for genome-wide methylation by MBD-seq. The analysis revealed 20 methylated genes, notably *Eif2B5*, that regulates myelin production. Loss of myelin with accompanying musculoskeletal pain is a major symptom of Gulf War illness. Our work demonstrates multiple genes were methylated by exposure to OPs and glucocorticoids. These genes point to biochemical mechanisms that may be targets for therapeutic intervention.

## Introduction

In 1991, a 42-nation coalition led by the United States, initiated combat against Iraq because of its invasion of Kuwait (Englehardt, 2019). Nearly one million personnel were involved and of those who participated in combat actions, between 25 and 35 percent became sick with a multi-symptom malaise now called Gulf War illness (GWI). Symptoms include gastrointestinal, respiratory, fatigue and cognitive problems. (Fappiano & Baraniuk, 2020). One of the hypotheses concerning the cause of the syndrome is exposure to organophosphates (OP), including sarin and chlorpyrifos, combined with increased circulating glucocorticoids as might be expected in the stress of combat. Moreover, for many of the afflicted veterans, the symptoms have persisted for more than 30 years since the conflict ended (Petry et al., 2024). Earlier research from our group presented evidence for genetic contributions to individual differences in susceptibility (Jones et al., 2020; Xu et al., 2020; Gao et al., 2020). In that series of experiments, we identified two possible candidate genes underlying individual differences in susceptibility to the exposures experienced by the troops in theater. That many of the afflicted veterans experience symptoms more than thirty years after cessation of the conflict poses a different, but important feature of GWI. The question is what is the reason these individuals are sick for so long? One highly likely possibility is the exposure modified the expression potential of one or more genes. Accordingly, we performed small studies to examine the possible methylation of genes, thus reducing their expression potential. The first of these involved the acute effects of a sarin surrogate coupled with a glucocorticoid (Ashbrook et al., 2018) in a single inbred mouse strain, C57BL/6J one of the foundation strains for the genetic reference family of recombinant inbred mouse strains, the BXD group (Peirce et al., 2004). The second study involved the two foundation strains for the BXD group, C57BL/6J and DBA/2J (Mozhui et al., 2023). In this study, we performed the same treatments, exposure to corticosterone followed by treatment with a sarin surrogate but harvesting the hippocampus for analysis of genome-wide methylation 12 weeks after the sarin surrogate treatment, to model the years-long persistent nature of the disorder in ill veterans. Here we report the results from a study that expanded the number of strains and time after treatment.

## Methods

### Mice and treatment

Male and female mice from11 BXD strains age 60-65 days of age were the subjects for this study. All subjects received corticosterone in their drinking water (20mg% w/v) for seven days. On the eighth day, the animals received 4.0 mg kg^-1^ i.p. diisopropylfluorophosphate (DFP), a sarin surrogate. Over the next 43 weeks and for the first 12 weeks on alternate weeks, the animals received corticosterone as above in their drinking water. On the 42^nd^ week, the animals received corticosterone in their drinking water. This was done to prime the neuroinflammatory effect of the original corticosterone/DFP treatment. Glucocorticoids are generally regarded as having anti-inflammatory effects; however, they can also show pro-inflammatory effects, (Frank et al., 2010; Bolshakov et al., 2021). One week following the last corticosterone treatment, the mice were euthanized, and the hippocampus was harvested. The DNA was extracted from the tissues and prepared for MBD-seq to identify genome-wide methylation and RNA extracted for analysis.

### Tissue harvest and sample preparation

Genomic DNA was extracted from the hippocampus using the Quick-DNA/RNA Miniprep Plus kit (Zymo Research, Irvine, CA, United States) and checked for purity and quantity using a NanoDrop spectrophotometer (ThermoFisher Scientific, Waltham, MA, United States), and a Qubit™ fluorometer and the dsDNA BR (Broad Range) Assay kit (Invitrogen). Affinity based CpG enrichment was done using the Invitrogen MethylMiner Methylated DNA Enrichment Kit (ThermoFisher Scientific, Waltham, MA, United States), which relies on the methyl-CpG binding domain protein 2 (MBD2) protein to capture DNA fragments containing methyl-CpGs. MBD2 preferentially binds to methylated CpGs, and this depletes the DNA sample of DNA regions without CpGs, and enriches for methyl-CpGs (Aberg et al., 2018; Aberg et al., 2020). First, 1 µg of DNA in 110 μL low TE (tris-EDTA) buffer was sheared to ∼150 bp fragments using a Covaris S2 ultrasonicator (Covaris, Woburn, MA, United States). Sonication settings were the same as described in Sandoval-Sierra (Sandoval-Sierra et al., 2020) with cycle/burst of 1 for 10 cycles of 60 s, duty cycle of 10%, and intensity of 5.0. DNA fragment size and quality were assessed using the Agilent Bioanalyzer 2100 (Agilent, Santa Clara, CA, United States). MBD-capture reaction was done according to the standard manufacturer’s protocol, followed by a single step elution with 2 M NaCl solution. The enriched DNA was then reconcentrated by ethanol precipitation, and the final concentration of methylated-CpG enriched DNA ranged from 0.17 to 2.1 ηg per μl (0.87 ± 0.39).

### Sequencing and initial data processing

Sequencing was performed to 40 million reads per sample (150 paired-end) on Illumina NovaSeq 6000 (Illumina, San Diego, CA, United States).

### Alignment to the reference genome

Mus musculus (mouse) reference genome (GRCm38) and gene model annotation files were downloaded from the Ensembl genome browser (https://useast.ensembl.org/). Indices of the reference genome were built using STAR v2.5.0a (Dobin et al., 2013) and paired-end clean reads aligned to the reference genome.

For quantification of DNA methylation, the binary alignment map files were loaded to the MEDIPS R package (version 1.44.0) (Lienhard et al., 2014) using the MEDIPS.createSet function with the following parameters: uniq = 1, extend = 150, ws = 150, shift = 0. This divided the mm10 mouse genome into 150 bp non-overlapping windows, and reads were counted for each bin. The MEDIPS. couplingVector function was used to compute the local CpG density (coupling factor or CF), and the read counts were normalized to the CF using the function MEDIPS.meth. To retain only 150 bp bins that had sufficient coverage for reliable quantification and statistical analyses, we implemented the following filters: 1) bins with no CpGs (CF = 0) and mean read counts ≤1 were excluded, resulting in 5,724,879 bins; 2) these were loaded to the EdgeR R package (version 3.34.1) (Robinson et al., 2010), and further filtered on the basis of counts per million (CPM) to retain only reads with more than 1 CPM in 2 or more libraries. This resulted in 210,191 CpG regions that were then normalized by the library size using the calcNormFactors function. RPKM values were then extracted using the parameters gene length = 150, log = TRUE, and the CpG regions were annotated using HOMER software (Heinz et al., 2010).

## Results

### CORT+DFP on DNA methylation

To define site specific changes induced by the long-term exposure to CORT+DFP, we performed an epigenome-wide association study (EWAS) to compare between the treatment and control groups. For the first test (EWAS 1), we examined the main effect of treatment, with sex and body weight as cofactor, and adjusted for the top 3 PCs as proxies of genetic heterogeneity and other sources of variance. At a nominal unadjusted p=0.0001, 34 CpG regions were altered in methylation by treatment (Data S1). The strongest main effect of treatment was on the promoter methylation of the *Eif2b5* gene (Fig 1a). This region showed no difference between the sexes and was significantly decreased in methylation by CORT+DFP (Fig 1b). Analysis of variance showed a significant treatment effect (F_1,65_=25.62, p<.001). The main effect for strain and strain X treatment interaction were not significant (F_10,65_=1.55, p<.2; F_10,65_=1.02, p<.4, respectively). The top 20 CpG regions associated with the CORT+DFP treatment are presented in Table 1. Many of the CpG regions were in intergenic sites, but also included CpGs located in the introns or exons of *Ctif, Cdh6*, etc.

**Table 1.** Methylated genes identified for Main effect of CORT+DFP treatment and sex by treatment interaction.

| GeneSymbol | Bin | Bin Bp | annotation | EWAS 1<sup>1</sup> |  | EWAS 2<sup>1</sup> |  |
| --- | --- | --- | --- | --- | --- | --- | --- |
|  |  |  |  | Treatment effect |  | Treatment effect |  |
|  |  |  |  | coef | p | coef | p |
| *Eif2b5* | 16 | 20501551 | Promoter (2-3kb) | 0.33 | 1.5E-07 | 0.35 | 3.5E-05 |
| *Ctif* | 18 | 75624151 | Exon (exon 3 of 12) | -0.35 | 5.0E-06 | -0.34 | 1.3E-03 |
| *2700062C07Rik* | 18 | 24484351 | Distal Intergenic | 0.30 | 8.2E-06 | 0.39 | 2.8E-05 |
| *Or2aj6* | 16 | 19650151 | Distal Intergenic | 0.30 | 1.2E-05 | 0.36 | 1.4E-04 |
| *Ctif* | 18 | 75624001 | Exon (exon 2 of 2) | -0.33 | 1.8E-05 | -0.21 | 3.8E-02 |
| *2410021H03Rik* | 17 | 69325201 | Intron (intron 2 of 2) | -0.21 | 3.4E-05 | -0.16 | 2.2E-02 |
| *Nsun2* | 13 | 69283501 | Distal Intergenic | 0.25 | 5.1E-05 | 0.19 | 1.9E-02 |
| *Nlrp4e* | 7 | 23315701 | Intron (intron 1 of 8) | -0.26 | 5.7E-05 | -0.33 | 2.7E-04 |
| *Gfod1* | 13 | 43327351 | Distal Intergenic | -0.29 | 5.9E-05 | -0.13 | 1.6E-01 |
| *Hlcs* | 16 | 94243501 | Intron (intron 1 of 6) | 0.29 | 6.2E-05 | 0.20 | 3.4E-02 |
| *Dph3* | 14 | 32112151 | Distal Intergenic | 0.29 | 6.2E-05 | 0.31 | 2.2E-03 |
| *Cdh6* | 15 | 13034251 | Exon (exon 12 of 12) | 0.27 | 6.8E-05 | 0.33 | 3.2E-04 |
| *Armh1* | 4 | 117353251 | Intron (intron 2 of 14) | 0.25 | 7.0E-05 | 0.27 | 2.2E-03 |
| *Zc3h12c* | 9 | 52416301 | Exon (exon 2 of 2) | 0.27 | 7.5E-05 | 0.35 | 2.1E-04 |
| *Stag3* | 5 | 138321601 | Distal Intergenic | -0.30 | 8.0E-05 | -0.33 | 1.8E-03 |
| *Maml3* | 3 | 51988651 | Exon (exon 2 of 2) | -0.23 | 8.8E-05 | -0.33 | 5.6E-05 |
| *Lrp6* | 6 | 134587651 | Distal Intergenic | 0.31 | 9.1E-05 | 0.34 | 1.9E-03 |
| *Sult3a2* | 10 | 33747001 | Distal Intergenic | 0.23 | 9.1E-05 | 0.30 | 2.6E-04 |
| *Kif26a* | 12 | 112277851 | Distal Intergenic | 0.28 | 9.4E-05 | 0.23 | 2.1E-02 |
| *Sphk1* | 11 | 116543851 | Exon (exon 1 of 9) | 0.23 | 9.9E-05 | 0.24 | 2.5E-03 |

<sup>1</sup>EWAS 1 tests for main effect of treatment with body weight, sex, and PCs as covariates; EWAS 2 same as EWAS 1 and includes sex-by-treatment interaction.

**Figure 1.**
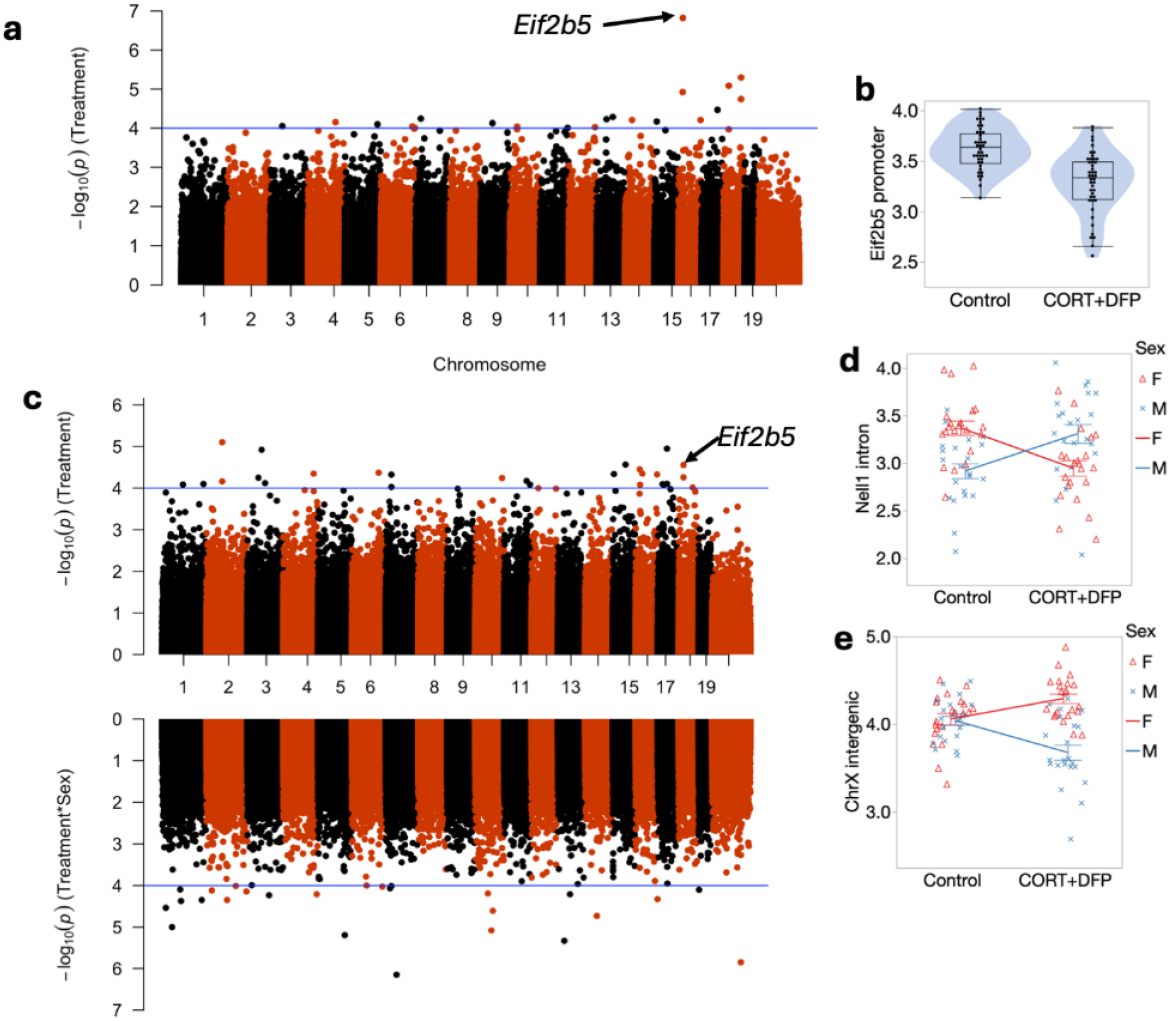
Manhattan plot of MBD-seq indices of genome-wide methylation. a) main effect of CORT+DFP; b) Effect of CORT+DFP on expression of the promoter region of *Eif2B5*; c) Manhattan plot following addition of sex X treatment; d) sex differences in methylation of *Eif2B5*; e) sex differences in treatment in an intergenic region on X chromosome.

We followed up the main EWAS with a regression that include the sex-by-treatment interaction (EWAS 2). The addition of this additional parameter reduced the associations for the main effect of treatment (Fig 1c). However, as a sensitivity test, the association with treatment for most of the CpG regions shown in Table 1 remained consistent at p<0.001. This also revealed CpGs that had evidence of sex-by-treatment interactions. The strongest interaction effect was for a CpG region in the intron of *Nell1* (alternatively, a promoter of the *Nell1os*). While this region shows no difference by treatment when all samples are pooled, but when stratified by sex, females show a reduction in methylation, while males show an increase in methylation with treatment (Fig 1d). An intergenic region on chromosome X showed a contrasting effect on treatment between the two sexes (Fig 1e). The top 20 CpG regions with treatment-sex interaction are in Table 2. None of these were associated with treatment based on EWAS 1.

### Relating to strain variation

As illustrated in Fig 2, many of the CpGs show significant heterogeneity by genetic background, and the top hits from the EWAS shown in Fig 1a are the ones that are robust to such heterogeneity.

**Figure 2.**
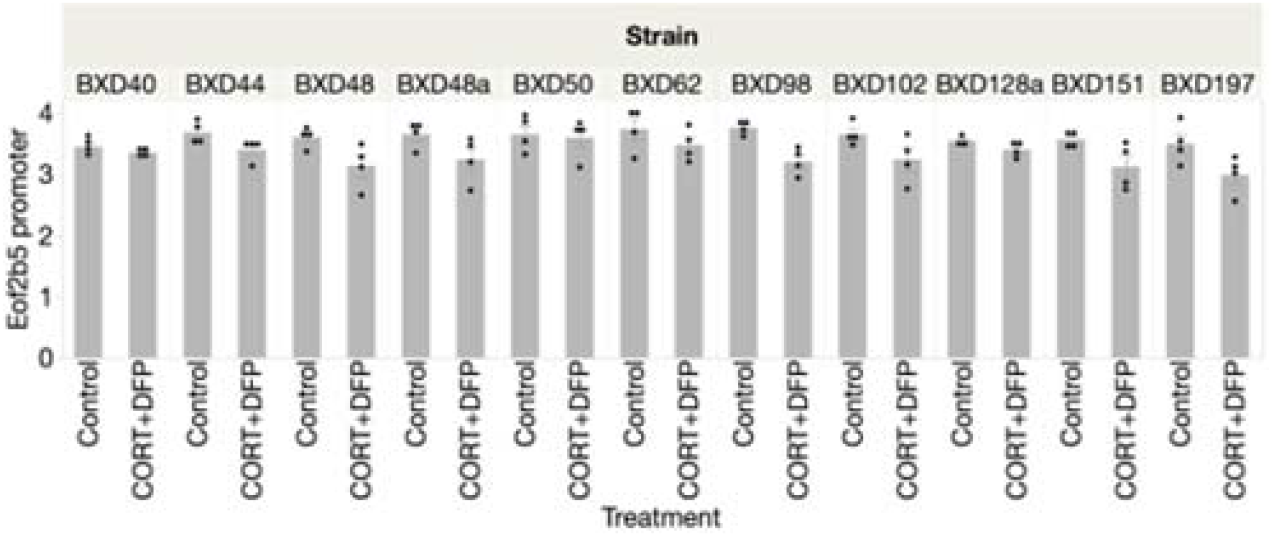
Strain distribution of methylation of *Eif2B5* 43 weeks following treatment with corticosterone and DFP

**Figure 3.**
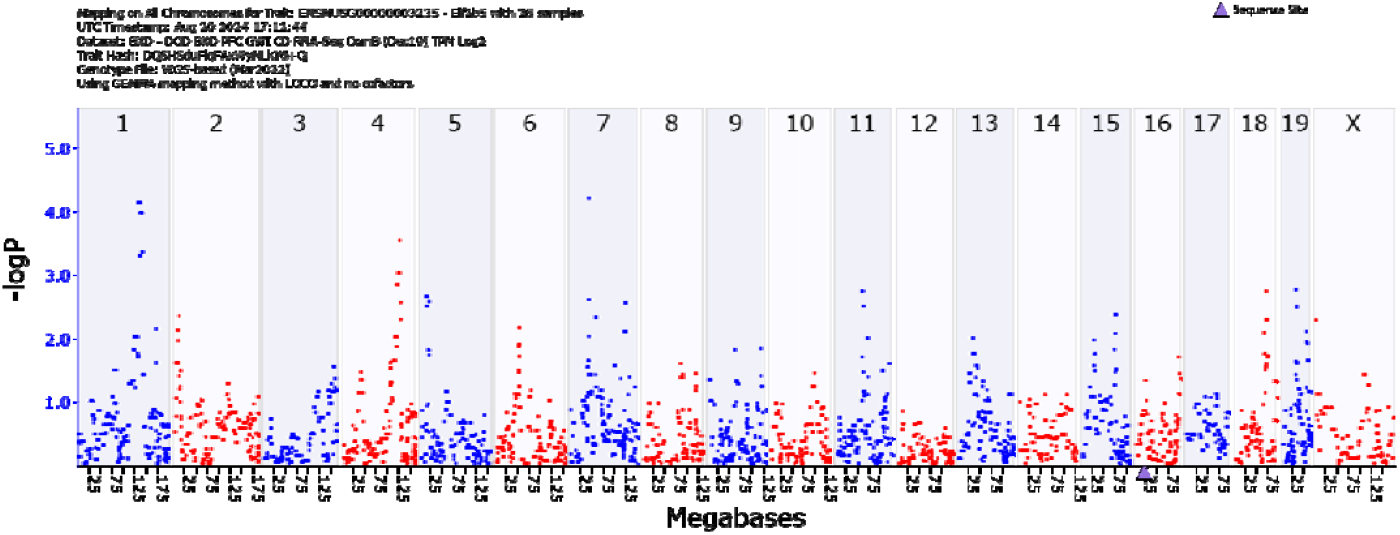
GEMMA quantitative trait analysis of *Eif2B5* gene expression. Note the significant association on Chromosome 1. Search for candidate genes in the interval reveals *Mapkapk2 (Mk2)*.

**Figure 4.**
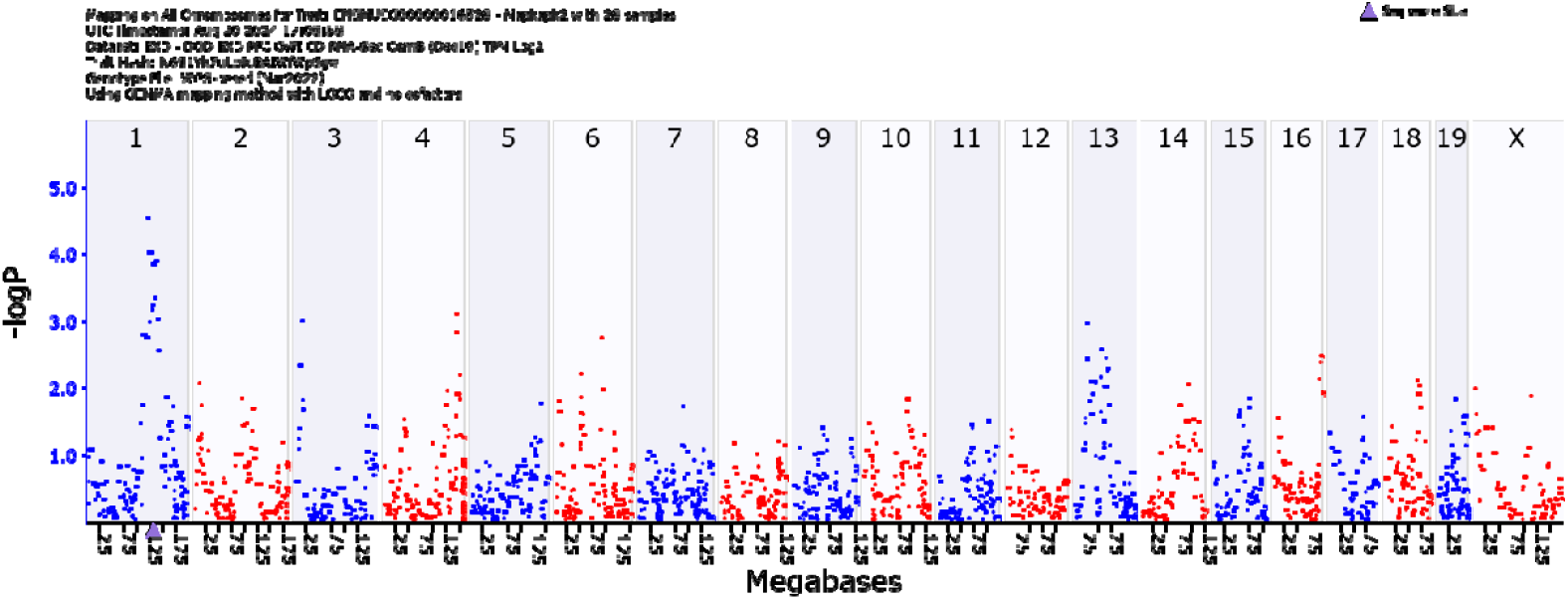
GEMMA quantitative trait analysis of *Mapkapk2* gene expression. Note the significant association on Chromosome 1. This indicates that the gene is *cis-*regulated.

Analysis of the MBDseq results showed significant associations between CORT+DFP for the following genes; *Eif2b5, Or2aj6, Nsun, Nlrp4e*, and *Cdh6*. The strongest signal was for *Eif2b5* (Figure 1a). The strain distribution of effect is presented in Figure 2.

## Discussion

Here, we extend the findings of our previous small study involving just the parental strains of the BXD group and for 84 instead of 300 days Mozhui et al., 2023). The small study was for proof of concept, and we identified four genes that had been differentially methylated by strain. The genes are *Ttll7*, which regulates glutamate as a neurotransmitter, *Akr1c14*, associated with the immune microenvironment, *Slc44a4*, a high-affinity, sodium-dependent choline transporter found in peripheral tissues and a choline supplier to other cells for membrane maintenance. *Rusc2* is associated with cognition. The present study expanded the number of BXD strains to 11 and extended the time from exposure to DFP to 300 days. The timeframe approximates the developmental age of Gulf War veterans who are now in their early to late 50s. The current study expands the number of methylated candidate genes, including Eif*2b5, Or2aj6, Nsun, Nlrp4e, Cdh6. Eif2b5* is associated with myelination and GWI includes neurological problems involving deficits in myelin. *Or2aj6*, facilitates translation, *Nsun* is a methyltransferase, *Nlrp4e* regulates interferon 1 and participates in inflammation. *Cdh6*, cahedrin 6, is associated with gliomas. Its expression is positively associated with malignancy and negatively associated with prognosis. This is an important player in GWI as those so afflicted have increased risk for glioma. Of particular interest is our finding that *Eif2b5* is regulated by *Mapkapk2*, a gene that is involved in cellular stress response, including production of the proinflammatory cytokines, *Tnfa* and *IL6*, among several other cellular processes.

As many of those afflicted with GWI are still experiencing debilitating symptoms, the research emphasis, now that probable etiological factors have been identified, the emphasis needs to be placed on treatments. One of the primary symptoms of chronic GWI is peripheral pain, originating from compromised central nervous system myelin (Van Riper et al., 2017). Vanishing white matter (VWM) disease is a fatal disorder but not related to the white matter compromise seen in GWI. This disease, triggered by stress, is seen primarily in children (Abbink et al., 2019). A recent study (Wong et al., 2019) reported that a small molecule compound, 2BAct, reversed the loss of myelin in a mouse model of VWM. This compound stimulates *Eif2B5* expression. This or similar compounds may have promise for treating the chronic debilitating symptoms of GWI.

## Conclusion

Here, we have shown a promising gene and biochemical mechanism that underlies chronic symptoms of GWI and the possibility of treating these symptoms. Because the GWI population is aging, time is the important factor in prosecuting the discovery of therapies.

## Acknowledgements

The authors thank Daming Zhuang for expert technical assistance Major support by USPHS grant R02ES031656

